# The bladder microbiome, metabolome, cytokines, and phenotypes in patients with systemic lupus erythematosus

**DOI:** 10.1101/2022.01.13.476281

**Authors:** Fengping Liu, Jingjie Du, Qixiao Zhai, Jialin Hu, Aaron W. Miller, Tianli Ren, Yangkun Feng, Peng Jiang, Lei Hu, Jiayi Sheng, Chaoqun Gu, Ren Yan, Longxian Lv, Alan J. Wolfe, Ninghan Feng

## Abstract

**Background and aims:** Emerging studies reveal a unique bacterial community in the human bladder, with alteration of composition associated to disease states. Systemic lupus erythematosus (SLE) is a complex autoimmune disease that is characterized by frequent impairment of the kidney. Here, we explored the bladder microbiome, metabolome, and cytokine profiles in SLE patients, as well as correlations between microbiome and metabolome, cytokines, and disease profiles.

**Methods and materials:** We recruited a cohort of 50 SLE patients and 50 individually matched asymptomatic controls. We used transurethral catheterization to collect urine samples, 16S rRNA gene sequencing to profile bladder microbiomes, and LC-MS/MS to perform untargeted metabolomic profiling.

**Results:** Compared to controls, SLE patients possessed a unique bladder microbial community and increased alpha diversity. These differences were accompanied by differences in urinary metabolomes, cytokines, and patients’ disease profiles. The SLE-enriched genera, including *Bacteroides*, were positively correlated with several SLE-enriched metabolites, including olopatadine. The SLE-depleted genera, such as *Pseudomonas*, were negatively correlated to SLE-depleted cytokines, including IL-8. Alteration of the bladder microbiome was associated with disease profile. For example, the genera *Megamonas* and *Phocaeicola* were negatively correlated with serum complement C3, and *Streptococcus* was positively correlated with IgG.

**Conclusions:** Our present study reveals associations between the bladder microbiome and the urinary metabolome, cytokines, and disease phenotypes. Our results could help identify biomarkers for SLE.

## Introduction

Systemic lupus erythematosus (SLE) is a complex autoimmune disease with a chronic relapsing-remitting course that can damage multiple organs and range from mild to life-threatening illness(1). The kidney is one of the most commonly impaired organs, and lupus nephritis (LN) has been reported in approximately 50% of SLE patients(2), 10-30% of LN patients progress to kidney failure that requires kidney replacement therapy(2), and the mortality rate within 5 years of onset directly attributed to kidney disease is 5-25% of patients with proliferative LN(2). Currently, no cure for SLE has been found and the pathogenesis of SLE is currently poorly understood.

Alteration of microbial compositions in the gut, oral mucosa or tegument has been reported to be associated with SLE disease manifestations(3–15). Specifically, SLE patients exhibit compositional alterations to the gut microbiome, characterized by lower bacterial diversity(3, 4, 6–8, 14), decreased Firmicutes/Bacteroidetes (F/B) ratio(5, 7, 14, 15), and increased abundance of Lactobacillaceae(6, 8, 11). Similarly, SLE patients exhibit reduced microbial diversity and altered microbial community in their gums and skin(10, 12). However, the association of SLE on microbial communities and metabolic output in the bladder and other urogenital niches has not been studied.

Once considered sterile, the bladder is now known to possess microbial communities (bladder microbiome) in individuals with and without urinary tract infections (16). Furthermore, disruption (dysbiosis) of the bladder microbiome is associated with urinary tract disorders, especially urgency urinary incontinence and urinary tract infection(17). However, most studies of bladder microbiome involve only US participants; only a few reports involve Chinese participants(18, 19).

Immune response and metabolic output can bridge the gap between the microbiome and SLE phenotypes. Urine is often used to assess metabolic status of the body(20). For example, Yan and co-workers found 23 metabolites dramatically increased in SLE patients compared to healthy controls, including valine, cysteine, and uracil(21). Also, as renal impairment is one of the most serious manifestations of SLE and urine cytokines derived directly from the diseased kidney accumulate in the urine, the level of inflammatory factors in urine may be used as an indicator of chronic inflammation and disease progression. Brugos and co-workers found that IL-1 and TNF-α were elevated significantly in the urine of patients without renal disease, while IFN-γ was elevated in the urine of LN patients(22). However, the relationship between immunity, metabolism, and microbiome in the bladder of SLE patients is unclear.

Given that SLE often affects the kidney, and LN is a common manifestation that leads to irreversible renal impairment, we hypothesize that the bladder microbiomes of individuals with and without SLE differ, and the differences correlate with specific urinary metabolites and cytokines, along with patients’ clinical profiles. To test this hypothesis, we analyzed urine obtained by transurethral catheterization from participants with and without SLE. We also compared the gut and vaginal microbiomes of a subset of SLE patients to their bladder microbiome to determine whether these adjacent microbiomes might influence the composition of the bladder microbiome.

## Methods and materials

### Patients and controls recruitment

50 SLE patients, who fulfilled at least 4 of the American College of Rheumatology Criteria for the diagnosis of SLE(23), and 50 sex-, age-, BMI-, and co-morbid disease-matched controls were consecutively recruited from Wuxi Second Affiliated Hospital of Nanjing Medical University (**Fig. S1**). The inclusion and exclusion criteria are described in **File. S1**. Disease activity was measured using SLEDAI score(24). All participants signed their informed consent before sample collection. The study was executed in accordance with the Ethical Committee of the hospital (ref. 201805).

### Urine sample collection

Urine samples were collected through a urinary catheter. Before insertion of the catheter, 5% iodophor was applied to sterilize the genital and perineal areas. The collected urine was separated into four portions, which were used for detecting or measuring the bladder microbiome, metabolome, creatinine levels, and cytokines. Fecal and vaginal samples were collected before the collection of urine samples. Fecal material was collected in a sterile container by the patient, and 30 mg was immediately placed in a sterile container. Vaginal samples were collected by the nurse using a sterile swab. All urine, feces and vaginal samples were placed in sterile, DNA- and enzyme-free centrifuge tubes, and immediately stored at −80°C until use.

### DNA extraction and bioinformatics analysis

30 mL urine samples were processed for sequencing as described in **File. S2.** The DNA extraction from the vaginal and fecal samples were the same as urine samples. We used DADA2 (https://github.com/benjjneb/dada2) to process reads derived from 16S rRNA V3-V4 region, including quality control (truncQ=8, maxN=0, maxEE=c(2,2)), dereplication, merging forward and reverse reads (trimOverhang=TRUE, minOverlap=5), and chimera removal (method=“consensus”) to obtain amplicon sequence variants (ASVs). To remove environmental contaminants, we manually removed ASVs whose reads did not exceed 5 times the maximum number of reads in the environmental controls. After decontamination, BLCA was applied to obtain taxonomic identities for the remaining ASVs(25). We only kept taxa with a confidence score above 60 for downstream analysis. Multivariate Association with Linear Modes (MaAslin) framework was used to adjust the effects of confounding factors.

### Urinary metabolome profiling and processing

Urinary metabolome profiling was performed using liquid chromatography tandem mass spectrometry, LC-MS/MS (ExionLC and TripleTOF 5600, SCIEX, Framingham, MA, USA) as previously described (see **File. S3**). In total, 6770 and 6078 peaks were detected in the positive and negative ionization modes, respectively. 6515 (positive mode) and 5970 (negative mode) metabolite features remained. The positive-mode and negative-mode features were then annotated using The Kyoto Encyclopedia of Genes and Genomes (KEGG, http://www.kegg.com/) and Human Metabolome Database (HMDB, http:www.hmdb.ca). The result was 1076 annotated metabolites (**Tab. S1**).

### Cytokines and creatinine detection

The Bio-Plex™ 200 System (Bio-Rad) and Bio-Plex Pro Human Cytokine 27-plex assay (Bio-Rad, California, USA) were used to detect urinary cytokines. Creatinine was detected using the human urine Elisa kit of creatinine (Hengyuan Biological Technology Co., Ltd, Shanghai, China).

### Disease profile measurement

Blood samples were collected on the day of urine sample collection. The immunoturbidimetric test was used to assess serum complement (C) and immunoglobulin antibodies (AU5421; Beckman Coulter, USA). Immunoblotting and immunofluorescence were used to detect serum autoantibodies. Erythrocyte sedimentation rate was determined by the Westergren method (XC-408 ESR Monitor; Mindray, China).

Lupus nephritis (LN) was defined as clinical and laboratory manifestations that meet ACR criteria(26). Systemic Lupus Erythematosus Disease Activity Index 2000 (SLEDAI-2K) was used to assess disease severity(24).

### Statistical analysis

For microbiome analysis, Bray-Curtis dissimilarity of microbial communities was calculated using ‘vegdist’ function with “bray” mode, permutational multivariate analysis of variance [PERMANOVA] was calculated using ‘adonis’ function, and Shannon’s H was calculated using ‘diversity’ function with “shannon” mode in R. For metabolome analysis, the concentration of urinary metabolites was adjusted for variability in urine dilution, using Cr as a normalization indicator. For the comparison of the metabolite intensity between SLE and controls, we performed statistical analysis using MetaboAnalyst 5.0 (https://www.metaboanalyst.ca). Metabolites with (1) variable importance in the projection (VIP) greater than 1, (2) fold change greater than 2 or less than 0.5 and (3) *P*-value less than 0.05, were then log2 transformed and subjected to linear model analysis to control for confounding factors, including nutrient intake (Binary logistic regression model analysis, SPSS 24.0). For cytokine analysis, Cr was used to normalize urinary cytokine concentration.

Pearson’s Chi-square or Fisher’s exact tests were used with categorical variables; Student’s *t* test was used on normalized continuous variables and Wilcoxon rank-sum test on non-normal continuous variables. The *P*-value was adjusted for multiple comparisons using the Benjamini-Hochberg (BH) false discovery rate (FDR).

## Results

### Demographics

We assessed the bladder microbiome, metabolite profile, and cytokine profile of a total of 50 SLE patients and 50 sex-, age-, BMI-, and comorbid disease-matched asymptomatic controls (Controls) (**Tab. 1**). Both cohorts were 88% (n=44) female, 12% (n=6) male. As expected, the SLE cohorts had lower serum concentrations of C3 and C4, but higher serum concentrations of IgG and uric acid; they also had more urinary white blood cells, red blood cells and leucocyte esterase (*P*<0.05 for all comparisons). Of the 50 SLE patients, 38 (76%) had LN (**Tab. S2**). The SLE cohort also had significantly elevated intake of calcium and zinc (**Tab. S3**); these were listed as confounding factors in the downstream analyses.

**Tab. 1.** Characteristics of controls and SLE patients.

| Parameters | Value for cohort (n <sup>a</sup>) <sup>b</sup> or statistic |  | P value <sup>c</sup> |
| --- | --- | --- | --- |
|  | Controls (n = 50) | SLE (n = 50) |  |
| Female sex, n (%) | 44 (88.00) | 44 (88.00) | 1.000 |
| Age (yrs) | 49.22 ± 15.19 | 45.00 ± 14.61 | 0.160 |
| Body-mass index (kg/m<sup>2</sup>) | 23.98 ± 3.23 | 23.99 ± 2.65 | 0.992 |
| Duration of SLE (yrs) | NA | 7.58 ± 5.97 | NA |
| History of smoking, n (%) | 0 (0.00) | 0 (0.00) | 1.000 |
| History of drinking, n (%) | 1 (2.00) | 0 (0.00) | 1.000 |
| **Comorbidity** |  |  |  |
| Diabetes, n (%) | 5 (10.00) | 5 (10.00) | 1.000 |
| Hypertension, n (%) | 10 (20.00) | 10 (20.00) | 1.000 |
| **Immunological features** |  |  |  |
| Complement 3 (g/L) | 1.20 ± 0.64 | 0.76 ± 0.23 | < 0.001 |
| Complement 4 (g/L) | 0.30 ± 0.28 | 0.16 ± 0.06 | < 0.001 |
| Ig A (g/L) | 2.59 ± 0.85 | 2.60 ± 1.08 | 0.930 |
| Ig G (g/L) | 10.98 ± 1.88 | 15.97 ± 7.86 | < 0.001 |
| Ig M (g/L) | 1.17 ± 0.51 | 1.01 ± 0.51 | 0.105 |
| ESR (mm/hr) <sup>d</sup> | NA | 30.02 ± 22.56 | NA |
| **Renal function** |  |  |  |
| Serum creatinine (μmol/L) | 55.23 ± 10.75 | 65.79 ± 40.62 | 0.087 |
| Blood urea nitrogen (mmol/L) | 5.73 ± 5.99 | 5.46 ± 2.11 | 0.766 |
| Serum uric acid (umol/L) | 268.04 ± 72.56 | 318.86 ± 103.66 | 0.006 |
| Estimated glomerular filtration rate (mL/min/1.73 m<sup>2</sup>) | 114.40 ± 20.08 | 108.84 ± 38.26 | 0.374 |
| Urinary creatinine (mg/dL) | 1.37 ± 0.19 | 1.42 ± 0.25 | 0.268 |
| **Urine analysis** |  |  |  |
| White blood cells (/uL) | 2.22 ± 6.38 | 9.32 ± 22.40 | < 0.001 |
| Red blood cells (/uL) | 0.61 ± 3.03 | 28.00 ± 131.45 | < 0.001 |
| Nitrites positive, n (%) | 1 (2.00) | 2 (4.00) | 1.000 |
| Leucocyte esterase, n (%) | 0 (0) | 12 (24.00) | < 0.001 |
| **LN** |  |  |  |
| LN, n (%) | NA | 38 (76.00) | NA |
| Duration of LN (yrs) | NA | 5.30 ± 3.60 | NA |
| **SLEDAI** |  |  |  |
| Score | NA | 16.16 ± 24.31 | NA |
| Low SLEDAI [<8; n(%)] |  | 8 (16.00) | NA |
| High SLEDAI [≥8; n(%)] |  | 42 (84.00) | NA |

<sup>a</sup> n, number of subjects;
<sup>b</sup> Mean ± SD or n (%);
<sup>c</sup> Pearson Chi-square or Fisher's exact test was used with categorical variables; Student's t test on normalized continuous variables and Wilcoxon rank-sum test was used on un-normalized continuous variables.
<sup>d</sup> ESR detection threshold was 15 mm/hr.
Abbreviations: ESR, erythrocyte sedimentation rate; Ig, immunoglobulin; LN, lupus nephritis; NA, not applicable;
SLEDAI, systemic lupus erythematosus disease activity index.

### Bladder microbiome is altered in SLE patients

To test whether the bladder microbiome differs between SLE patients and asymptomatic controls, we first assessed the microbial community structure using all microbial species present within the bladder microbiome of each sample. Principal Coordinate Analysis (PCoA) of Bray-Curtis dissimilarities revealed differential clustering between the control and SLE cohorts (**Fig. 1A**; R^2^=0.128, *P_(adj)_*=0.001), reflecting a dysbiotic urobiome in SLE patients. To determine whether the bacterial community was affected by medication usage, we separated the SLE patients into subgroups based on their dosages of hydroxychloroquine and prednisone and performed PCoA; we found no differences between/among the subgroups (**Fig. S2A and S2B**; R^2^=0.013, *P_(adj)_*=0.886; R^2^=0.020, *P_(adj)_*=0.428). The disruption of microbial composition between the controls and SLE cohort was highlighted by the observation that species diversity (as measured by the Shannon’s H Index) was significantly elevated in the SLE cohort (**Fig. 1B**; *P_(adj)_*<0.05;), likely due to increased evenness (**Fig. S3A**; *P_(adj)_* <0.05;), as there was no significant difference in species richness (**Fig. S3B**).

**Fig. 1.**
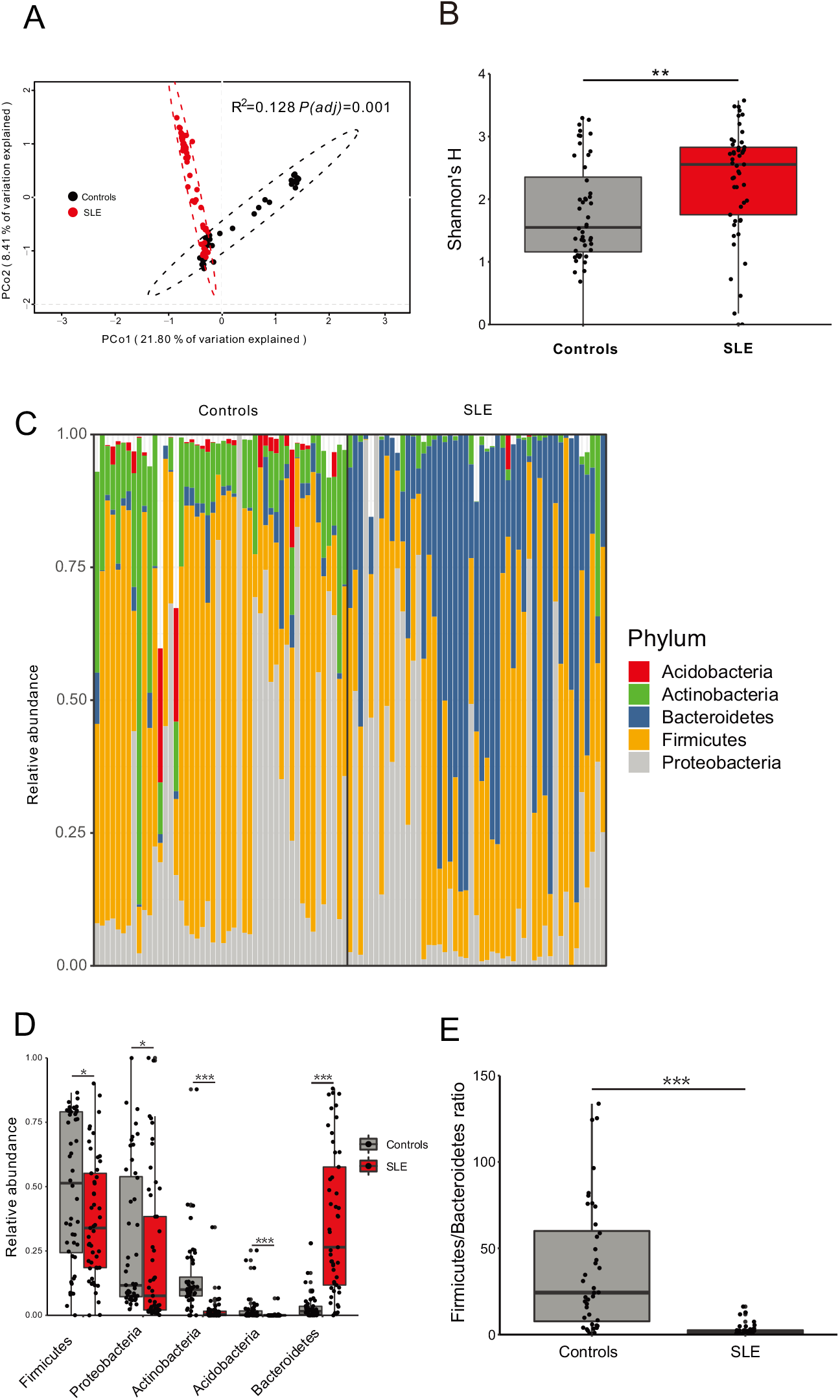
Bacterial composition, diversity and phylum difference between controls and SLE cohort. A. PCoA based on Bray-Curtis distances at species level showed different microbial compositions between groups of SLE patients and controls. The 95% confidence ellipse is drawn for each group. Permutational multivariate analysis of variance (PERMANOVA) was performed for statistical comparisons of samples in the two cohorts. *P* value was adjusted by the Benjamini and Hochberg false discovery rate (FDR). B. Bacterial diversity measured by Shannon index was calculated at the bacterial species level. Wilcoxon rank-sum test was performed and adjusted by Benjamini and Hochberg false discovery rate (FDR). ** indicates *P_(adj)_*<0.01. C. Microbial profile at the phylum level. Only phyla with more than 1% average relative abundances in all samples are shown. D. Bacterial phyla that were differentially abundant between controls and SLE patients. *P* value was calculated using Wilcoxon rank-sum test and adjusted by Benjamini and Hochberg false discovery rate (FDR). * and *** indicate *P_(adj)_*<0.05 and *P_(adj)_*<0.001, respectively. E. Firmicutes/Bacteroidetes ratio differed in controls and SLE patients. *P* value was calculated using Wilcoxon rank-sum test and adjusted by Benjamini and Hochberg false discovery rate (FDR). *** indicates *P_(adj)_*<0.001.

Since we observed a clear difference in diversity, we assessed taxonomic signatures at the phylum level. The relative abundances of the 5 most abundant phyla (>1% relative abundance) differed significantly between the control and SLE cohorts (**Fig. 1C**; *P_(adj)_*<0.05). The phyla Firmicutes, Proteobacteria, Acidobacteria and Actinobacteria were significantly more abundant in controls, whereas Bacteroidetes was significantly more abundant in the SLE cohort (**Fig. 1D**; *P_(adj)_*<0.05). The Firmicutes/Bacteroidetes ratio was reduced significantly in SLE patients (**Fig. 1E**; *P_(adj)_*<0.05), consistent with previous studies of the gut microbiome of SLE patients(5, 27).

At the genus level, PCoA based on the Bray-Curtis Dissimilarity Index also revealed differential clustering of bladder microbiomes from controls relative to SLE patients (**Fig. S4A**; R^2^=0.153, *P_(adj)_*=0.001). The 15 most abundant genera (>1% relative abundances) are displayed in **Fig. S4B**. Among them, seven genera, (*Staphylococcus, Rothia, Streptococcus, Haemophillus, Sphingomonas, Gardnerella* and *Pseudomonas*) were significantly more abundant in controls (**Fig. 2A**; *P_(adj)_*<0.05), especially *Staphylococcus*, which often predominated. In contrast, 5 genera (*Alistipes, Bacteroides, Phocaeicola, Phascolarctobacterium* and *Megamonas*), were significantly more abundant in the SLE cohort (**Fig. 2B**; *P_(adj)_*<0.05). However, when we adjusted for the confounding factors, such as nutrient intake and medication usage, using MaAslin analysis, we found that *Alistipes* and *Blautia wexlerae* were affected by calcium intake, and *B. wexlerae* was also affected by prednisone use (**Tab. S4**; *P_(adj)_* <0.001); thus, *Alistipes* and *B. wexlerae* were removed from the downstream interaction analysis. The bacterial species >0.5% relative abundances are displayed in **Fig. S5**; 8 species were significantly more abundant in controls, especially *S. aureus* (**Fig. 2C**), whereas 11 species were significantly more abundant in the SLE cohorts (**Fig. 2D**). To investigate the potential for the use of urinary microbial profiles to discriminate SLE patients from controls, we performed a backward stepwise selection model to identify bacterial genera/species with optimal model fitting. This model identified 3 genera (*Bacteroides, Rothia* and *Sphingomonas)* and 4 species (*Phocaeicola vulgatus, Rothia aeria, Asticcacaulis excentricus* and *Vicinamibacter silvestris*) that could discriminate controls from SLE with AUC values of 93.16% (**Fig. 2E**) and 98.63% (**Fig. 2F**), respectively.

**Fig. 2.**
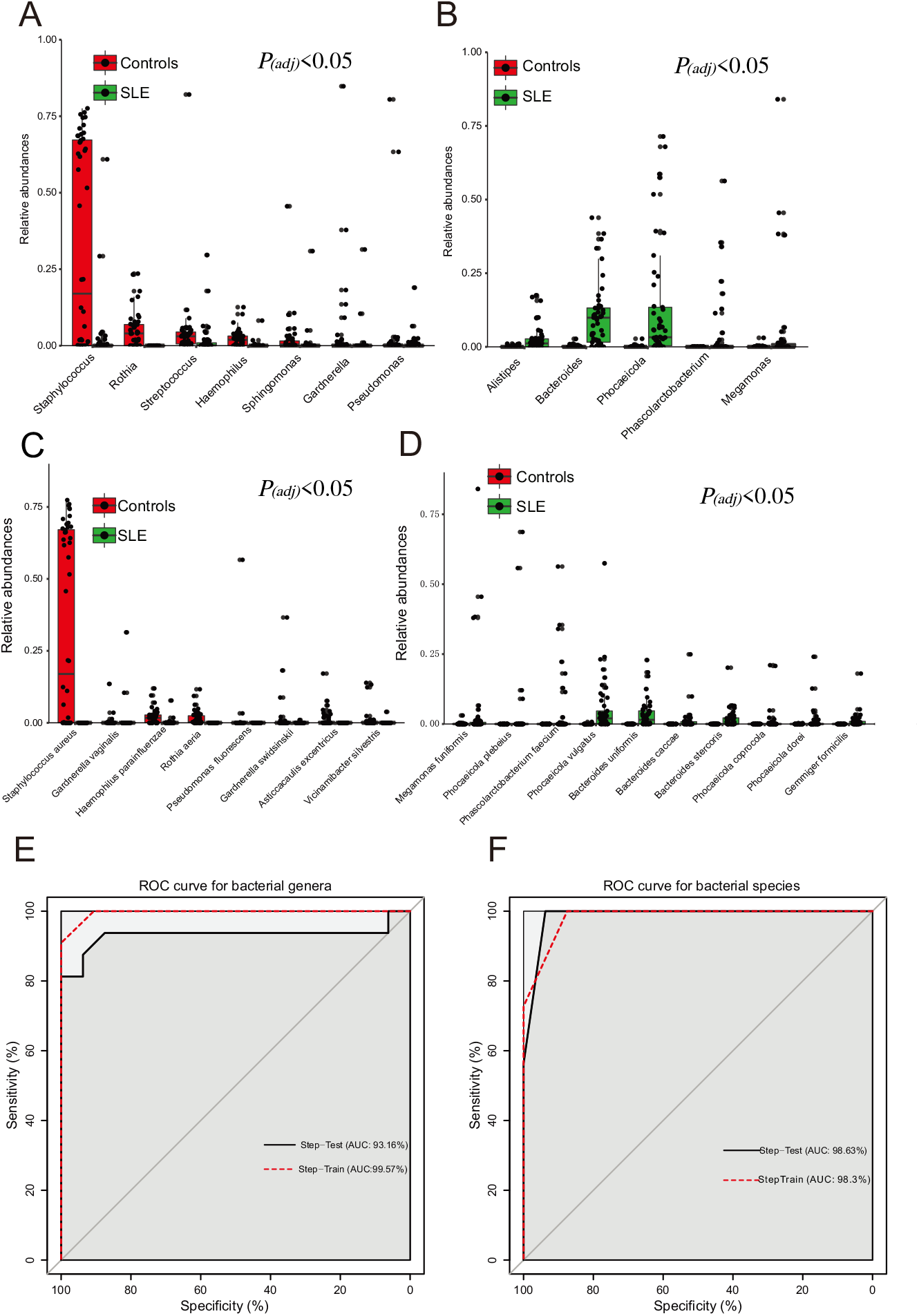
Bacterial genera and species are different between controls and SLE cohort. A. Bacterial genera that were more abundant in controls compared to SLE patients (*P_(adj)_*<0.05). *P* value was calculated using Wilcoxon rank-sum test and adjusted by Benjamini and Hochberg false discovery rate (FDR). B. Bacterial genera that were less abundant in controls compared to SLE patients (*P_(adj)_*<0.05). *P* value was calculated using Wilcoxon rank-sum test and adjusted by Benjamini and Hochberg false discovery rate (FDR). C. Bacterial species that were more abundant in controls compared to SLE patients (*P_(adj)_*<0.05). *P* value was calculated using Wilcoxon rank-sum test and adjusted by Benjamini and Hochberg false discovery rate (FDR). D. Bacterial species that were less abundant in controls compared to SLE patients (*P_(adj)_*<0.05). *P* value was calculated using Wilcoxon rank-sum test and adjusted by Benjamini and FDR. E. ROC curve for bacterial genera. A backward stepwise selection model to identify bacterial genera with optimal model fitting. This model identified 3 genera (*Bacteroides, Rothia* and *Sphingomonas*) that could discriminate SLE from controls with AUC values of 93.16%. F. ROC curve for bacterial species. A backward stepwise selection model to identify bacterial species with optimal model fitting.

Next, we divided SLE patients into lupus nephritis (LN) and non-LN subgroups (**Tab. S5**); the LN and non-LN cohorts matched demographically, except for age. Based on PCoA, the bladder microbiome of these subgroups did not differ (R_2_=0.022, *P_(adj)_*=0.377), but both differed from controls (**Fig. S6A**; R^2^=0.072, *P_(adj)_*=0.002; R^2^=0.128, *P_(adj)_*=0.002, respectively). As measured by Shannon’s H Index, microbial diversity also did not differ between these two subgroups, but the SLE patients with LN had more microbial diversity than controls (**Fig. S6B**; *P_(adj)_*0.05).

Finally, as a pilot analysis, we compared the bladder microbiome with the vaginal and gut microbiomes using only the subset (N=15) of SLE patients who provided all 3 sample types. PCoA based on species showed that the SLE microbiome of the bladder differed from those of the vagina and gut (**Fig. S7A**; R^2^=0.105, *P_(adj)_*=0.001). By Bray-Curtis Dissimilarity Index, the SLE bladder microbiome more closely resembled the vaginal microbiome than it did the gut microbiome (**Fig. S7B;** *P_(adj)_*<0.001, and *P_(adj)_*<0.01, respectively). However, the predominant species in the bladder microbiome were dissimilar to those of both the gut and vagina (**Fig. S7C**).

### Urinary metabolome is altered in SLE patients

To test whether the urinary metabolome differs between SLE patients and controls, we performed untargeted metabolomics on the urine samples. Based on Principal Component Analysis (PCA) using all 1076 metabolites detected (**Tab. S1**), the metabolic composition of SLE patients differed significantly from controls (**Fig. 3A**; R^2^=0.650, *P_(adj)_*=0.001). Partial least squares discriminant analysis (PLS-DA) yielded similar results (**Fig. 3B**; *P*<0.001). Also, we tested the effect of medication usage on the metabolome and found no differences between/among the dosage subgroups (**Fig. S8A and 8B;** R^2^=0.004, *P_(adj)_*=0.420; R^2^=0.058, *P_(adj)_*=0.892). These results suggested that the metabolome differed between controls and SLE patients, and the difference was not due to medication.

**Fig. 3.**
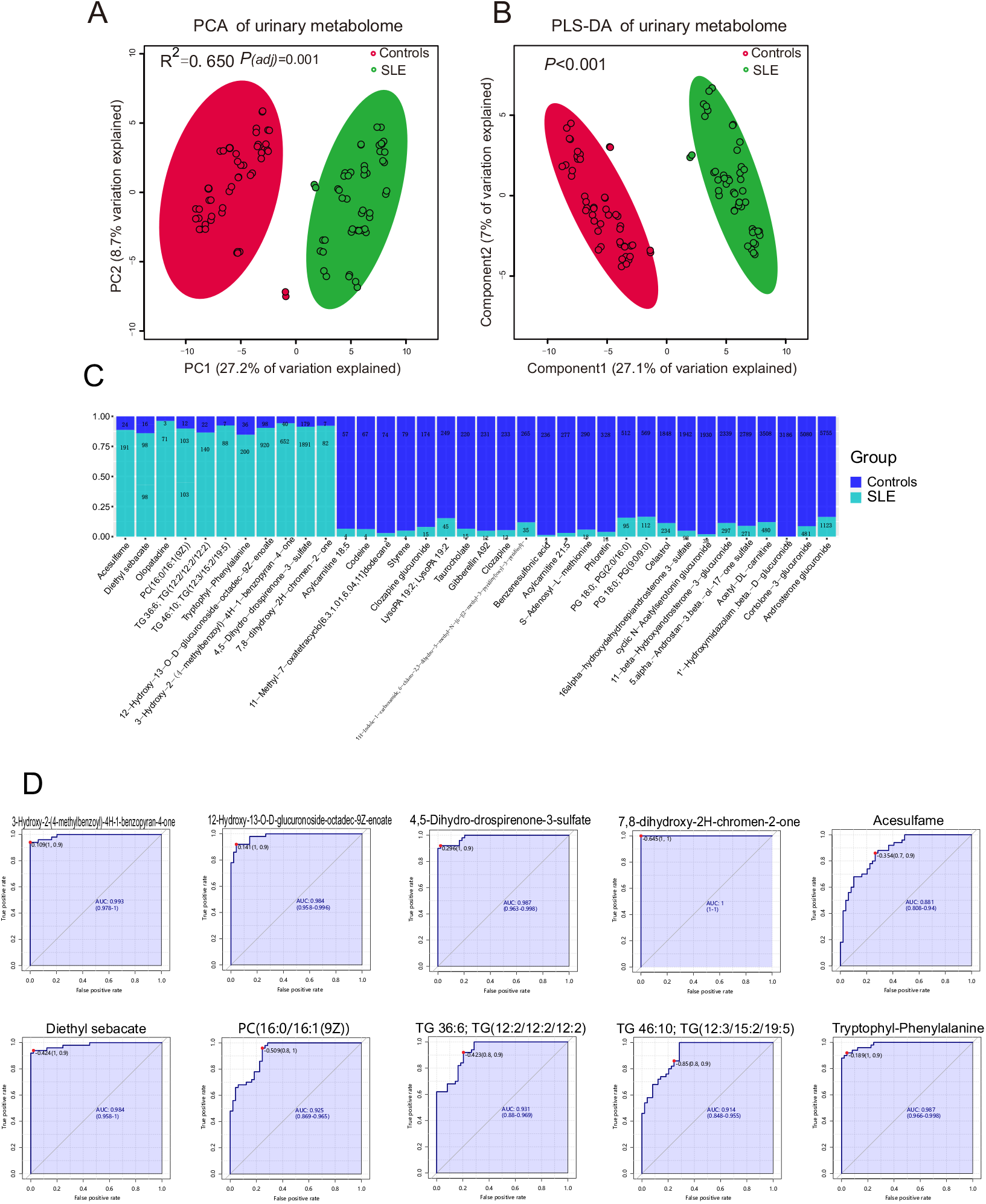
Urinary metabolome differed in SLE patients. A. Separation of urinary metabolome between patients with SLE and controls, revealed by principal component analysis (PCA). The explained variances are shown in brackets. Permutational multivariate analysis of variance (PERMANOVA) was performed for statistical comparisons of samples in two cohorts. *P* value was adjusted using the Benjamini and Hochberg false discovery rate. The 95% confidence ellipse is drawn for each group. B. Partial Least Square-Discriminant Analysis (PLS-DA) plot. Scores plot between the selected PCs. The explained variances are shown in brackets. PERMANOVA was used to test statistical comparisons of ions in SLE and control cohorts. C. The metabolites showing significant difference between the control and SLE cohorts. The metabolites described in the graph met the following criteria: *P_(adj)_* < 0.05 in Wilcoxon rank-sum test; variable importance in projection (PLS-DA; VIP>1) in Partial Least Square-Discriminant Analysis; and fold change (FC) > 2 or < 0.5. D. Receiver operating characteristic curve (ROC) curve for validation of metabolomic classification of control and SLE patients The sensitivity is on the y-axis, and the specificity is on the x-axis. The area-under-the-curve (AUC) is in blue.

Of the 1076 annotated metabolites, 120 metabolites were significantly more abundant in the SLE cohort, whereas 124 were significantly less abundant (**Fig. S9A;** *P*<0.05). Among the top 25 most abundant metabolites in the heatmap, 13 were visually less abundant in SLE group (**Fig. S9B**). After adjusting for confounding factors, including nutrient intake, 38 metabolites with variable importance in the projection (VIP) greater than 1 and with fold change less than 0.5 or greater than 2 differed significantly between controls and SLE patients (**Tab. S6-9**, *P*<0.05, **Fig. 3C**). To determine whether these 38 metabolites were affected by medication usage, we compared the metabolites according to dosage, and found no difference (**Tab. S10-11**; *P_(adj)_*>0.05). Since urinary hydroxychloroquine and desethylchloroquine are metabolites of the medication hydroxychloroquine(28), binary regression analysis was used to determine whether they were affected by hydroxychloroquine intake. It showed that hydroxychloroquine intake was a confounding factor of urinary hydroxychloroquine and desethylchloroquine (**Tab. S12**); thus, they were removed in the downstream analysis. To look for potential biomarkers that could distinguish SLE from controls, classical ROC curve analysis (including logistic regression analysis with selected variables to get the modeling results and compare the performance using the accuracy/performance plots i.e. area under the curve, specificity, and sensitivity) was used to evaluate the performance of single metabolites. From this analysis, 10 metabolites had an AUC value above 0.85, indicating they could be biomarkers of SLE (**Fig. 3D**).

Like the bladder microbiome, the metabolic composition of the LN and non-LN subgroups did not differ, but each differed significantly from the composition of controls (**Fig. S10;** Controls vs LN, R^2^=0.255,*p_(adj)_*=0.002; Controls vs nonLN, R^2^=0.173, *P_(adj)_*=0.002; LN vs nonLN, R^2^=0.010, *P_(adj)_*=0.909). However, when we compared the metabolic differences between controls and LN, 427 metabolites were differentially abundant (*P_(adj)_*<0.05); 185/427 had fold change greater than 2 or less than 0.5 and 121/185 metabolites with VIP greater than 1 (**Tab. S13-15**). When the control and non-LN cohorts were compared, 239 metabolites were differentially abundant (*P_(adj)_*<0.05), 132/239 metabolites with fold change greater than 2 or less than 0.5, and 101/132 metabolites with VIP greater than 1 (**Tab. S16-18**).

### Bladder microbiome was associated with urinary metabolome

Bladder microbiome and urinary metabolome correlated robustly across all subjects (**Fig. S11**, M^2^=0.906, *P*=0.001). To determine specific associations between the bacterial genera and metabolites, we conducted a Spearman correlation analysis using the abundant bacterial genera (>1% relative abundances) and metabolites that differed between the SLE and control cohorts. Indeed, most of the SLE-enriched genera were positively correlated with most of the SLE-enriched metabolites, and most of the SLE-depleted genera were negatively correlated with most of the SLE-enriched metabolites (**Fig. 4 and Tab. S19;** |r|>0.3, *P*<0.05). For example, the SLE-enriched genera, such as *Bacteroides*, were positively correlated with SLE-enriched metabolites, such as the lipids and lipid-like molecules, including PC (16:0/16:1(9Z)). Notably, the SLE-enriched genera, including *Bacteroides*, were positively correlated with several SLE-enriched organohetercocyclic compounds, including olopatadine.

**Fig. 4.**
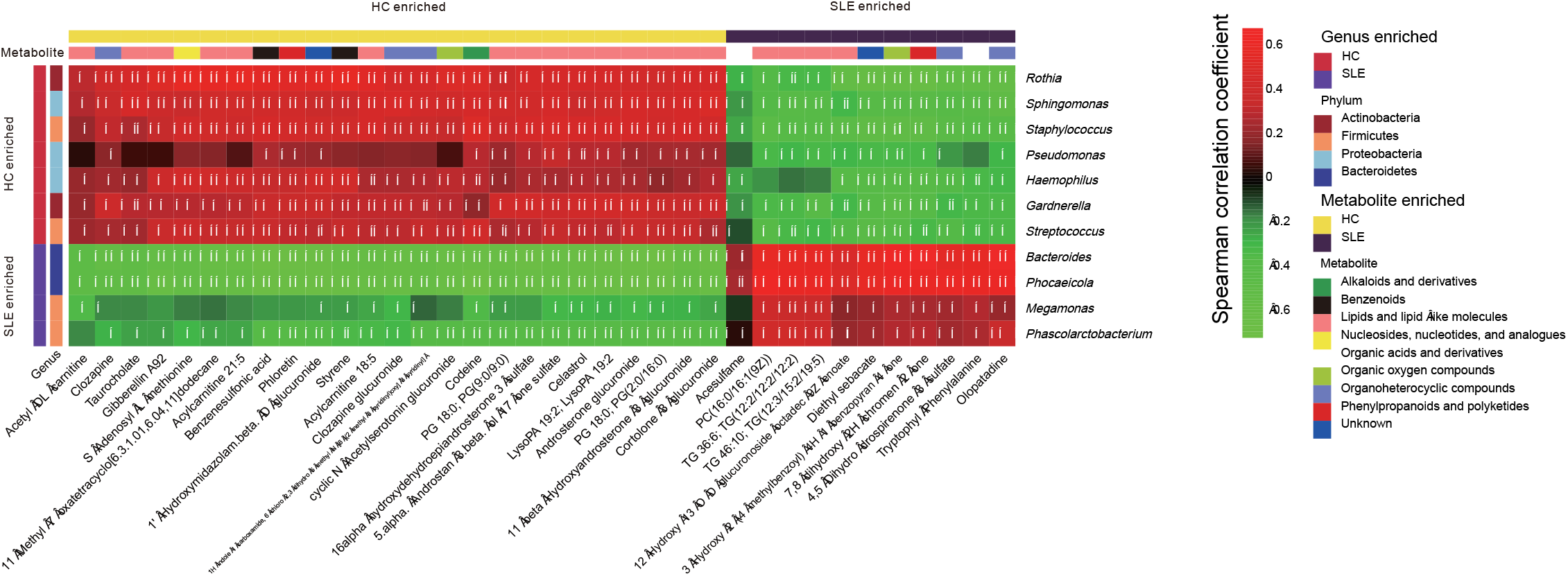
Bladder microbiome was associated with metabolites. The heatmap depicted the association between the taxa and metabolites that differ in SLE relative to controls. Spearman correlation analysis was performed on the abundant bacterial genera (>1% relative abundances) and metabolites that differed between the control and SLE cohorts. The correlation of two variables with values of |r|>0.3 and *P*<0.05 are displayed. *, **, and *** indicate *P*<0.05, *P*<0.01 and *P*<0.001, respectively.

### Urine cytokines were altered in SLE patients

We next tested whether urinary cytokines differed between SLE patients and controls; 26 of the 27 cytokines assessed were identified. Among them, 10 cytokines differed in concentration between SLE patients and controls, including 6 cytokines (Eotaxin, G-CSF, IL-8, IL17, IP-10, and MIP-1b) significantly more abundant in SLE patients and 4 cytokines (IL-2, IL-5 IL-12 and IL-13) significantly less abundant in SLE patients (**Fig. 5A;** *P_(adj)_*<0.05). However, when we compared the cytokines among the controls, LN, and non-LN SLE patients, 12 cytokines, including IL-8, differed significantly between controls and LN SLE patients, and no cytokines differed significantly between controls and non-LN SLE patients (**Fig. S12;** *P_(adj)_*<0.05).

**Fig. 5.**
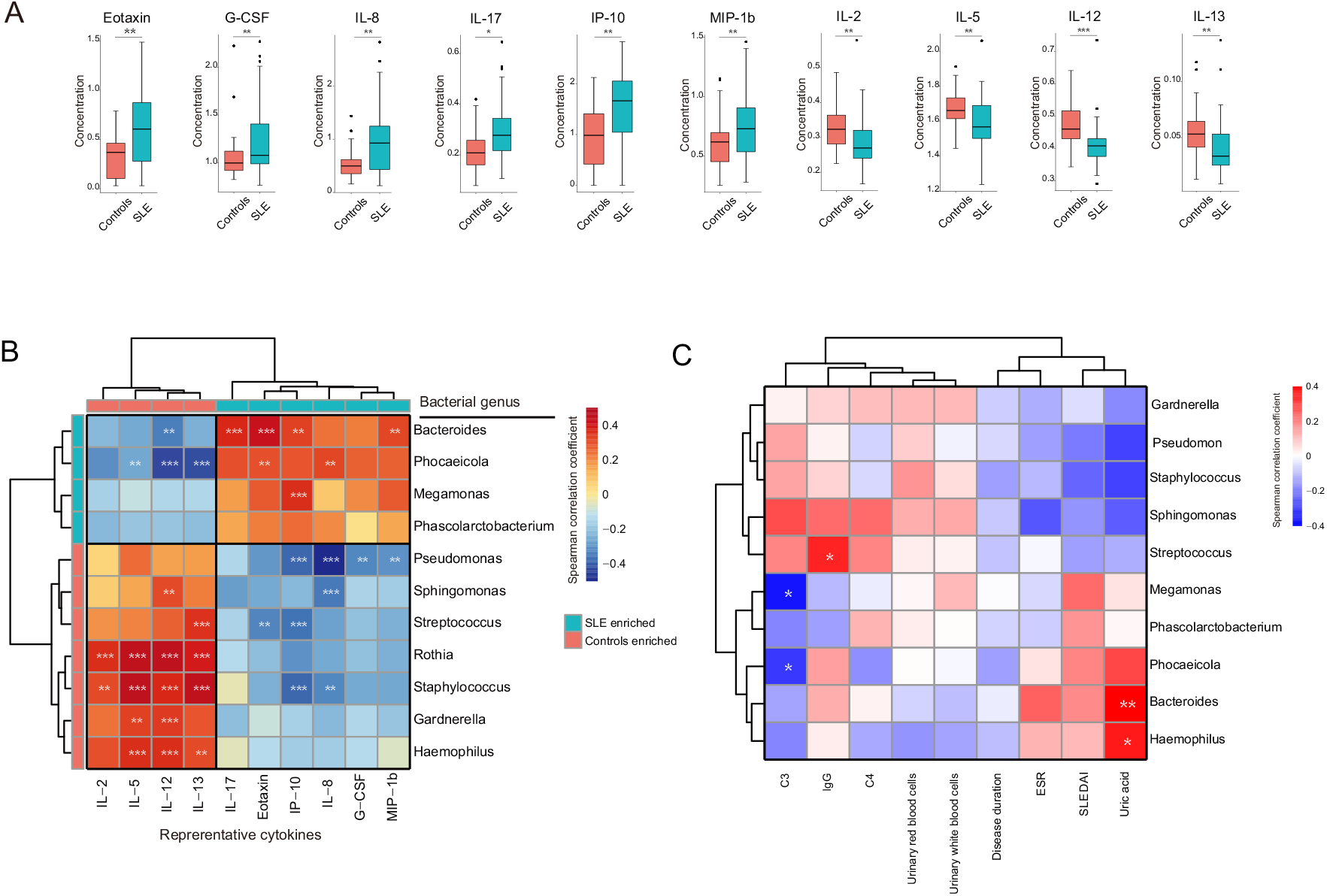
Urinary cytokines and profiles in SLE were associated with bladder microbiome. A. Urinary cytokines increased and decreased in SLE cohort compared to controls. *P* value was calculated using Wilcoxon rank-sum test and adjusted by Benjamini and Hochberg false discovery rate. B. Spearman correlation analysis was performed on the most abundant bacterial genera (>1% relative abundances) and cytokines that differed between the controls and SLE cohorts. The correlation of two variables with values of |r|>0.3 and *P*<0.05 are displayed. *, **, and *** indicate *P*<0.05, *P*<0.01 and *P*<0.001, respectively. C. Spearman correlation analysis was performed on the abundant bacterial genera (>1% relative abundances) and disease profiles of SLE patients. The correlation of two variables with values of |r|>0.3 and *P*<0.05 are displayed. *, **, and *** indicate *P*<0.05, *P*<0.01 and *P*<0.001, respectively.

### Bladder microbiome was correlated to urine cytokines

To examine the associations between differentially abundant bacterial genera and SLE-linked cytokines, we performed a correlation analysis (**Fig. 5B and Tab. S20**, |r|>0.3, *P*<0.05). Several of the SLE-enriched bacterial genera were positively associated with several SLE-enriched cytokines. In contrast, the SLE-depleted genera, such as *Pseudomonas*, were negatively correlated to SLE-deleted cytokines like IL-8.

### Bladder microbiome was associated with SLE-linked disease profiles

To look for associations between bladder microbiome and SLE-linked disease profiles, we performed a correlation analysis between the bacterial genera with the patients’ disease profiles. The disease profiles were as follows: the duration of SLE and LN, SLEDAI, immunological features, and urinary analysis outcomes that displayed significant differences between controls and SLE cohort or those only be detected in the SLE patients (**Tab. 1**). *Megamonas* and *Phocaeicola* were negatively correlated with serum complement C3, whereas IgG, more abundant in SLE patients, was positively correlated with *Streptococcus*. *Bacteroides* and *Haemophilus* were positively correlated to uric acid (**Fig. 5C and Tab. S21,** |r|>0.3, *P*<0.05).

## Discussion

This is the first integrated multi-omics analysis of bladder urine. It shows that the bladder microbiome and urinary metabolome profiles of SLE patients differs significantly from those of controls. It also shows that the microbiome profile correlates substantially with the urinary metabolome, urinary cytokines, and disease characteristic profiles.

First, we noticed that the bladder microbiome was associated with SLE. Compared with asymptomatic controls, the SLE bladder microbiome was greatly altered with increased microbial diversity and altered abundances of individual taxa at all tested taxonomic levels. Alterations to the microbiomes of SLE patients have been reported for human gut, oral cavity, and skin(3–15); however, the increased diversity of the SLE bladder microbiome was dissimilar to the findings of previous studies of the SLE gut microbiome, in which patients had lower bacterial diversity than controls(3, 4, 8, 11). The increased diversity of the bladder microbiome is quite substantial, considerably larger than patients with lower urinary tract symptoms(29, 30).

The bladder microbiome of asymptomatic controls, most of whom were women from eastern China (Jiangsu province), was most often predominated by *Staphylococcus*. This differs from the *Lactobacillus*-predominant bladder microbiome of asymptomatic controls from the upper midwestern U.S (Chicago area)(31, 32). Different genetic, ethnic, sociocultural, lifestyle, and dietary diversity may contribute to differences in the bladder microbiomes of North American and Chinese women. A large cohort study is needed to verify the differences and investigate the underlying factors that contribute to those differences.

The phylum Bacteroidetes, the genera *Bacteroides* and *Alistipes*, and the species *B. uniformis* were most evidently more common in SLE cohorts compared with controls, which resembles a previous study on the SLE gut microbiome(7). The stark contrast in Bacteroidetes between our SLE patients and asymptomatic controls warrants further investigation. *Staphylococcus* and *Pseudomonas* were less common in members of the SLE cohort. A previous bladder microbiome study showed that *Staphylococcus* was associated with urgency urinary incontinence(31). This was an unexpected result as these genera are considered pathogenic in SLE-associated infections(33). This is a striking result that should be investigated.

LN is a form of glomerulonephritis that constitutes one of the most severe organ manifestations in SLE patients. Previous human and animal gut microbiome studies demonstrated that LN is associated with microbiome composition(4, 34). Thus, we divided the patients into LN and non-LN subgroups. Although there was no significant difference in bacterial composition or abundance between LN and non-LN patients, microbial diversity differed between LN patients and controls, indicating there might be an interaction between bladder microbiome and inflammation in patients’ kidneys. However, the cohort sizes were too small and imbalanced to draw a strong conclusion.

The urinary metabolome also differed significantly in our study. Previous studies also showed that the urinary metabolome was altered in SLE and LN patients(35, 36), but those studies analyzed voided urine, which often contains post-urethral contamination(37). We assessed catheterized urine that avoided those contaminants(37), and identified several metabolites altered in SLE patients. For example, organohetercocyclic compounds, including olopatadine, were more abundant in SLE patients.

Like the microbiome and metabolome profiles, bladder cytokines also differed between SLE patients and controls. For example, IL-8 was significantly more abundant in SLE patients, consistent with previous cytokine studies of SLE patients(38, 39). As the kidney is affected by SLE and evaluation of urinary cytokines is reported to predict development of renal flares(40), we compared the urinary cytokines among controls and both LN and non-LN SLE patients. No cytokines differed between controls and non-LN SLE patients. In contrast, 10/12 of the cytokines that differed between controls and LN SLE patients also differed between controls and the entire cohort of SLE patients. The frequently confirmed LN-associated cytokines, such as IL-17(41), IP-10 and MCP-1(42), were only more abundant in the LN group compared to controls. These findings suggest that kidney damage in SLE patients may be responsible for altered bladder cytokine expression.

As integration of microbiome and metabolome data have potential for identifying microbial influence on host physiology through production, modification, and/or degradation of bioactive metabolites, we performed correlation analysis on the bladder microbiome and metabolome data. A major finding of the present study is the positive association between *Bacteroides* and organohetercocyclic compounds including olopatadine. Olopatadine is reported to be an antihistamine agent with inhibitory activities against chronic inflammation(43). *Bacteroidetes* also was positively correlated to PC (16:0/16:1(9Z)), which has been reported to have anti-inflammatory properties(44). Thus, the interaction between *Bacteroides* and organohetercocyclic compounds in the bladder might play a potential role in inhibiting inflammation-associated metabolites in SLE.

The microbiome plays a vital role in the regulation of host mucosal inflammation. To investigate interactions between bladder microbes and the inflammation response, we performed the integration analysis between the altered bacterial genus and urinary cytokines in patients. The SLE-enriched genus *Bacteroides* was positively correlated to the SLE-enriched cytokines, including IP-10 and IL-17, which are usually used as biomarkers to predict disease severity(41, 45). We also noticed that *Pseudomonas* was negatively correlated to IL-8, which can induce chemotaxis in target cells, and cause neutrophils and granulocytes to migrate toward the inflammation site(46). In our study, IL-8 was elevated only in LN patients and not non-LN patients, which fits with these IL-8 functions. Therefore, *Pseudomonas* might play a role in regulating immunity in bladder microbial community. The correlations between bladder microbiome and urinary cytokines indicate there may be a microbe-inflammation axis that should be explored.

It is known that complement deficiency is associated with SLE, predisposing these patients to infection(47). Indeed, the SLE patients in this study had decreased complement C3 and C4 levels and a negative correlation between C3 and SLE-enriched bacterial genera, including *Megamonas*. In contrast, *Megamonas* has been reported to be depleted in the gut of SLE patient and positively correlated to Th17(48). These observations warrant further investigation of *Megamonas* in SLE pathology. In addition, the SLE-depleted genus, *Streptococcus*, was positively correlated to immunoglobulin G (IgG), a major serum immunoglobulin principally responsible for elimination of pathogens and toxic antigens(49). Uric acid accumulation is common in SLE patients(50); it was elevated in our study, and positively correlated with *Bacteroides* and *Haemophilus*. Based on these findings, we hypothesize the possible existence of a host-microbe interaction in the human bladder that contributes to the SLE phenotype.

Individuals with SLE are regularly treated with immunosuppressives, which can cause serious adverse effects that severely compromise life quality. Therefore, there is an urgent need to control the disease process. Our present study demonstrated that the bladder microbiome of SLE patients is associated with their urinary metabolites, cytokines, and disease profiles, highlighting plausible disease-specific mechanisms for future investigation. In addition, we noticed a more distinct alteration of the microbiome, metabolome, and cytokines in LN patients compared to non-LN patients. Future mechanistic studies should focus on LN patients.

Our study has several limitations. Given frequent infection in SLE patients, it was hard to recruit volunteers who were willing to accept urinary catheterization (which is an invasive procedure); therefore, our sample size was small and entirely recruited at a single center with participants from eastern China. Moreover, as females are more susceptible to SLE, only a few males were recruited to this study. Thus, we could not compare sexual differences. A study with a larger sample size that includes more males and participants from different areas of China should be conducted in the future.

## Acknowledgments

We gratefully acknowledge the volunteers who participated in our study.

## Data accession

Raw data from 16S rRNA sequencing are available in the Sequence Read Archive (SRA) under BioProject ID PRJNA758740 (https://www.ncbi.nlm.nih.gov/bioproject/?term=PRJNA758740).

## Contributions

Conceptualization: Alan J. Wolfe, Longxian Lv, Ninghan Feng

Methodology: Fengping Liu, Tianli Ren, Qixiao Zhai,

Software: Fengping Liu, Jingjie Du, Ren Yan

Validation: Alan J. Wolfe

Writing: Alan J. Wolfe, Fengping Liu, Jingjie Du, Aaron W. Miller, Longxian Lv

Supervision: Alan J. Wolfe

Funding acquision: Ninghan Feng

Project administration: Fengping Liu, Jialing Hu, Lei Hu, Peng Jiang, Jiayi Sheng, Chaoqu Gu, Yangkun Feng

## Supplementary Figures

**Fig. S1** Overview of the study design.

**Fig. S2 Medication usages in patients didn’t affect bladder microbiome.**

A. PCoA based on Bray-Curtis distances at species level did not show different microbial compositions between SLE patients taking hydroxychloroquine dosages of 0.2 mg/d and those taking 0.4 mg/d. Permutational multivariate analysis of variance (PERMANOVA) was performed for statistical comparisons of samples using different levels of hydroxychloroquine. *P* value was adjusted by Benjamini and Hochberg false discovery rate.

B. PCoA based on Bray-Curtis distances at species level did not show different microbial compositions among SLE patients taking prednisone of 0 mg/d, 5 mg/d and those taking 10 mg/d. PERMANOVA was performed for statistical comparisons of samples using different levels of prednisone. *P* value was adjusted by Benjamini and Hochberg false discovery rate.

**Fig. S3 Bacterial evenness and richness of bacterial diversity.**

A. Comparison of bacterial species diversity indicator of evenness between control and SLE cohorts. Permutational multivariate analysis of variance (PERMANOVA) was performed for statistical comparisons of samples in two cohorts. *P* value was adjusted by Benjamini and Hochberg false discovery rate. ** indicates *P_(adj)_*<0.01.

B. Comparison of bacterial species diversity indicator of richness between control and SLE cohorts. PERMANOVA was performed for statistical comparisons of samples in two cohorts. *P* value was adjusted by Benjamini and Hochberg false discovery rate.

**Fig. S4 Bacterial community and composition at bacterial genus level.**

A. PCoA based on Bray Curtis distances at the genus level showed different microbial compositions between control and SLE cohorts. Permutational multivariate analysis of variance (PERMANOVA) was performed for statistical comparisons of samples in two cohorts. *P* value was adjusted by Benjamini and Hochberg false discovery rate.

B. Heatmap of bacterial genera. The 15 abundant genera (>1% relative abundances) were displayed.

**Fig. S5 Bacterial composition at bacterial species level.**

The heatmap displays the bacterial species >0.5% relative abundances.

**Fig. S6 Comparison of bladder microbiome among control, LN and non-LN cohorts.**

A. Bacterial community among controls, LN and non-LN SLE patients. PCoA based on Bray Curtis distances at species level was performed. Permutational multivariate analysis of variance (PERMANOVA) was performed for statistical comparisons of samples in two cohorts. *P* value was adjusted by Benjamini and Hochberg false discovery rate.

B. Bacterial diversity measured by Shannon index was calculated at the bacterial species level. Wilcoxon rank-sum test and adjusted by Benjamini and Hochberg false discovery rate (FDR). ** indicates *P_(adj)_*<0.01.

**Fig. S7 Comparison of microbiome in bladder, vagina, and gut in SLE patients.**

A. PCoA based on Bray-Curtis distances at species level showed different microbial compositions between the bladder, vagina, and gut. (PERMANOVA) was performed for statistical comparisons of samples in two cohorts. *P* value was adjusted by Benjamini and Hochberg false discovery rate.

B. Bray-Curtis dissimilarities of the different niches were calculated using the same SLE patient. Wilcoxon rank-sum test and adjusted by Benjamini and Hochberg false discovery rate (FDR). **, *** indicate *P_(adj)_*<0.01 and *P_(adj)_*<0.001, respectively.

C. Microbial profiles of the bladder, gut and vagina at species level. Bacterial species abundance with>0.05 % relative abundances are displayed. On the x axis, the numbers 1 to 15 represent SLE patients with bladder, gut and vaginal samples sequenced.

**Fig. S8 Medication usages in SLE patients didn’t affect urinary metabolome.**

A. Principal component analysis (PCA) was used to compare urinary metabolome between SLE patients taking hydroxychloroquine dosages of 0.2 mg/d and those taking 0.4 mg/d. The explained variances are shown in brackets. Permutational multivariate analysis of variance (PERMANOVA) was performed for statistical comparisons of samples in two cohorts. *P* value was adjusted using the Benjamini and Hochberg false discovery rate. The 95% confidence ellipse is drawn for each group.

B. Principal component analysis (PCA) was used to compare urinary metabolome among SLE patients taking prednisone of 0 mg/d, 5 mg/d, and 10 mg/d. Permutational multivariate analysis of variance (PERMANOVA) was performed for statistical comparisons of samples in two cohorts. *P* value was adjusted using the Benjamini and Hochberg false discovery rate. The 95% confidence ellipse is drawn for each group.

**Fig. S9 Metabolome comparison between control and SLE cohorts.**

A. Volcano plot. Volcano plot of differential metabolites classification of the control and SLE cohorts. Metabolites with FDR < 0.05 obtained by non-parametric tests and fold change (FC) >2 were identified as significantly different between the two cohorts. Colored plots indicate upward trend and downward trend of metabolites, and gray plots indicate that they are not statistically significant.

B. Clustering result shown as heatmap. Distance was measured using Euclidean, and clustering algorithm was calculated using Ward’s method.

**Fig. S10 Metabolome comparison among control, LN and non-LN SLE cohorts.**

Metabolome among controls, LN and non-LN SLE patients were compared using principal component analysis(PCA). Permutational multivariate analysis of variance (PERMANOVA) was performed for statistical comparisons of samples between groups. *P* value was adjusted by Benjamini and Hochberg false discovery rate. The 95% confidence ellipse is drawn for each group.

**Fig. S11 The relationship between bladder microbiome and urinary metabolome.**

Procrustes analysis analyzed the congruence of two-dimensional shapes produced from superimposition of principal component analyses from the datasets of microbiome and metabolome. Euclidian distances of eigenvalues for both the microbiome and metabolome using the Procrustes function in the vegan R package. Longer lines on Procrustes plots indicate more within-subject dissimilarity of the microbiome and metabolome. Significance value shown was calculated using the protest function from the vegan R package.

**Fig. S12 Urinary cytokines differed in LN SLE patients comparing to controls.**

Comparison of urinary cytokines between controls and LN SLE patients. *P* value was calculated using Wilcoxon rank-sum test and adjusted by Benjamini and Hochberg false discovery rate. *, **, *** indicate *P_(adj)_*<0.05, *P_(adj)_*<0.01 and *P_(adj)_*<0.001, respectively.

## Supplementary Tables

**Tab. S1 Annotable metabolites.**

Untargeted urinary metabolite profile was performed on liquid chromatography tandem mass spectrometry.

**Tab. S2** SLE patient’s characteristics.

Abbreviations: “-” and “+” represent negative and positive, respectively; ACA, anti-centromere antibodies; AHAs, Antihistone antibodies; AMA-M2, Anti-mitochondrial M2 antibody; ANAs, antinuclear antibodies; Anti-dsDNA, anti-double stranded DNA; Anti-NCS, anti-nucleosome antibodies; Anti-nRNP, anti-nRNP antibodies; Anti-PCNA, antibodies to the proliferating cell nuclear antigen; Anti-PM/Scl, anti-PM/Scl antibodies; Anti-Ro/SSA, anti-Ro/SSA antibodies; Anti-Ro52, anti-Ro52 antibodies; Anti-Scl-70, anti-Scl 70 antibodies; Anti-Sm, anti-Smith antibodies; Anti-SSB, anti-SSB antibodies; ASO, anti-streptolysin O; ESR, erythrocyte sedimentation rate; HCQ, hydroxychloroquin; IgG, immunoglobulin G; Meth, methylprednisolone; MTX, methotrexate; Pred, prednisone; RF, rheumatoid factor

**Tab. S3 Comparison of nutrient intake between controls and SLE.**

Student’s *t* test on normalized continuous variables and Wilcoxon rank-sum test was used on un-normalized continuous variables.

**Tab. S4 Bacterial taxonomy affected by food intake.**

MaAsLin (Microbiome Multivariable Associations with Linear Models) was used to adjust confounding factors, food intake, on bacteria showing significant difference at their abundance using Wilcox test (Galaxy Version 1.0.1). *P* value was adjusted using Benjamin Hochberg false discovery rate (FDR).

**Tab. S5 Demographics of LN and non-LN patients**

^a^ n, number of subjects;

^b^ Mean ± SD or n (%);

^c^ Pearson Chi-square or Fisher’s exact test was used with categorical variables; Wilcoxon rank-sum test was used on un-normalized continuous variables.

Abbreviations: LN, lupus nephritis; NA, not applicable; SLEDAI, systemic lupus erythematosus disease activity index

**Tab. S6 Metabolite comparison between using wilcox rank test (Controls and LN)**

Wilcoxon rank sum test was used on the metabolites. *P* value was adjusted using Benjamin Hochberg false discovery rate (FDR).

**Tab. S7 Metabolites showing Fold change>2 or <0.5 (Controls vs SLE)**

For paired fold change analysis, the algorithm first counts the total number of pairs with fold changes that are consistently above/below the specified FC threshold >2 or <0.5 for each variable.

**Tab. S8 Metabolites with VIP >1 (Controls vs SLE)**

VIP was calculated using PLS-DA analysis.

Abbreviation: VIP, Variable Importance in Projection

**Tab. S9 The effects of nutrient intake on metabolites showing significant different between controls and SLE**

Binary logistic regression model was used.

Abbreviations: B, coefficient value; SE, standard error; df, degrees of freedom; 95% CI, 95% confidence interval

**Tab. S10 Effects of hydroxychloroquine intake on metabolites**

Wilcoxon rank-sum test was used on the metabolites. *P* value was adjusted using Benjamin Hochberg false discovery rate (FDR).

**Tab. S11 Effects of prednisone intake on metabolites**

Wilcoxon rank-sum test was used on the metabolites. *P* value was adjusted using Benjamin Hochberg false discovery rate (FDR).

**Tab. S12 The effects of hydroxychloroquine intake on urinary hydroxychloroquine and Desethylchloroquine**

Binary logistic regression model was used.

Abbreviations: B, coefficient value; SE, standard error; df, degrees of freedom; 95%CI, 95% confidence interval

**Tab. S13 Metabolite comparison between controls and LN**

Wilcoxon rank-sum test was used on the metabolites. *P* value was adjusted using Benjamin Hochberg false discovery rate (FDR).

Abbreviation: LN, lupus nephritis

**Tab. S14 Metabolite comparison between controls and LN showing Fold change>2 or <0.5**

For paired fold change analysis, the algorithm first counts the total number of pairs with fold changes that are consistently above/below the specified FC threshold >2 or <0.5 for each variable.

**Tab. S15 Metabolites with VIP >1 (Controls vs LN)**

VIP was calculated using PLS-DA analysis.

Abbreviation: LN, lupus nephritis; VIP, Variable Importance in Projection

**Tab. S16 Metabolite comparison between controls and non-LN**

Wilcoxon rank-sum test was used on the metabolites. *P* value was adjusted using Benjamin Hochberg false discovery rate (FDR).

Abbreviation: non-LN, non-lupus nephritis

**Tab. S17 Metabolites showing Fold change>2 or <0.5 (controls vs non-LN)**

For paired fold change analysis, the algorithm first counts the total number of pairs with fold changes that are consistently above/below the specified FC threshold >2 or <0.5 for each variable. Abbreviation: non-LN, non-lupus nephritis

**Tab. S18 Metabolites with VIP >1 (Controls vs non-LN)**

VIP was calculated using PLS-DA analysis.

Abbreviation: LN, lupus nephritis; VIP, Variable Importance in Projection

**Tab. S19 Correlation between bacterial genus and metabolites that showed significant difference between controls and SLE**

Spearman correlation evaluated the linear relationship between bacterial genus and metabolite.

**Tab. S20 Correlation between bacterial genus and cytokines that showed significant difference between controls and SLE**

Spearman correlation evaluated the linear relationship between bacterial genus and cytokine.

**Tab. S21 Correlation between bacterial genus and disease profiles in SLE patients**

Spearman correlation evaluated the linear relationship between bacterial genus and disease profile.

